# Schizokinen siderophores in the methylotrophy model organism *Methylorubrum extorquens* AM1

**DOI:** 10.64898/2026.05.05.723055

**Authors:** Ignacio Sottorff

## Abstract

The facultative methylotroph model organism *Methylorubrum extorquens* AM1 is a known lanthanide user, which has shed light on the role of rare-earth metals in biochemistry. The characterization of a methanol dehydrogenase (MDH) protein which requires lanthanides as an enzymatic cofactor outlined the question of how these metals are acquired from the environment. It has been proposed that mesophilic organisms as *M. extorquens* AM1 can produce siderophore-like molecules, which chelate, transport and traffic rare-earth elements into the microbial cell. Therefore, we performed the bioinformatic and chemical investigation of *M. extorquens* AM1 by using genome mining, the CAS and arsenazo assay, molecular networking and chemical analytical techniques. Our results showed that indeed *Methylorubrum extorquens* AM1 harbored a gene cluster to produce metal chelators. The chemical analysis confirmed the production of the known hybrid hydroxamate-citrate siderophores schizokinen A and N-deoxyschizokinen A, which are very likely the side products of the transformation of schizokinen and N-deoxyschizokinen. The determination of the lanthanide chelation activity of the schizokinen siderophores series against three different lanthanides (La, Eu and Lu) showed no coordination activity, thus ruling out the involvement of schizokinen siderophores in rare-earth metal transport.

## INTRODUCTION

Organisms that use C_1_ reduced chemicals are called methylotrophs. The main chemicals for methylotrophy are methane and methanol (Chistoserdova and Kalyuzhnaya 2018). The use of C_1_ molecules can be obligate or facultative, depending on the organism physiological and ecological conditions (Chistoserdova et al. 2009). A model organism for the study of methylotrophy is *Methylorubrum extorquens* (Cotruvo et al. 2018), which has been isolated from the phyllosphere of *Arabidopsis thaliana* (Knief et al. 2010). *M. extorquens* is a Gram-negative, pink-pigmented bacterium that belongs to the phylum Proteobacteria and to the family *Methylobacteriaceae* (Garrity et al. 2005). Often, the strains that belong to the genera *Methylobacterium* and *Methylorubrum* are referred as pink pigmented facultative methylotroph, PPFM (Green and Ardley 2018).

*Methylorubrum extorquens* has been studied biochemically by its capacity to use methanol and lanthanides, being one of few organisms with a clear rare-earth biological use (Chistoserdova and Kalyuzhnaya 2018). In a lesser extent, it has also been analyzed for secondary metabolites production. To date, a handful of molecules have been characterized (**Fig. 1**), which can be associated to two different groups of secondary metabolites, acyl-homoserine lactones (Nieto Penalver et al. 2006) and toblerols. Homeserine lactones are commonly associated with the process of chemical communication among bacteria, particularly with quorum sensing (Schauder et al. 2001). Toblerols are a group Trans-AT polyketide synthetase (type I PKS) products which bear the uncommon cyclopropanol moiety (Ueoka et al. 2018b). Cyclopropanols are known to be methanol dehydrogenase (MDH) inhibitors (Frank Jr et al. 1989), as well as modulators of antibiosis in the bacterial genus *Methylorubrum* (Ueoka et al. 2018b).

**Figure 1.**
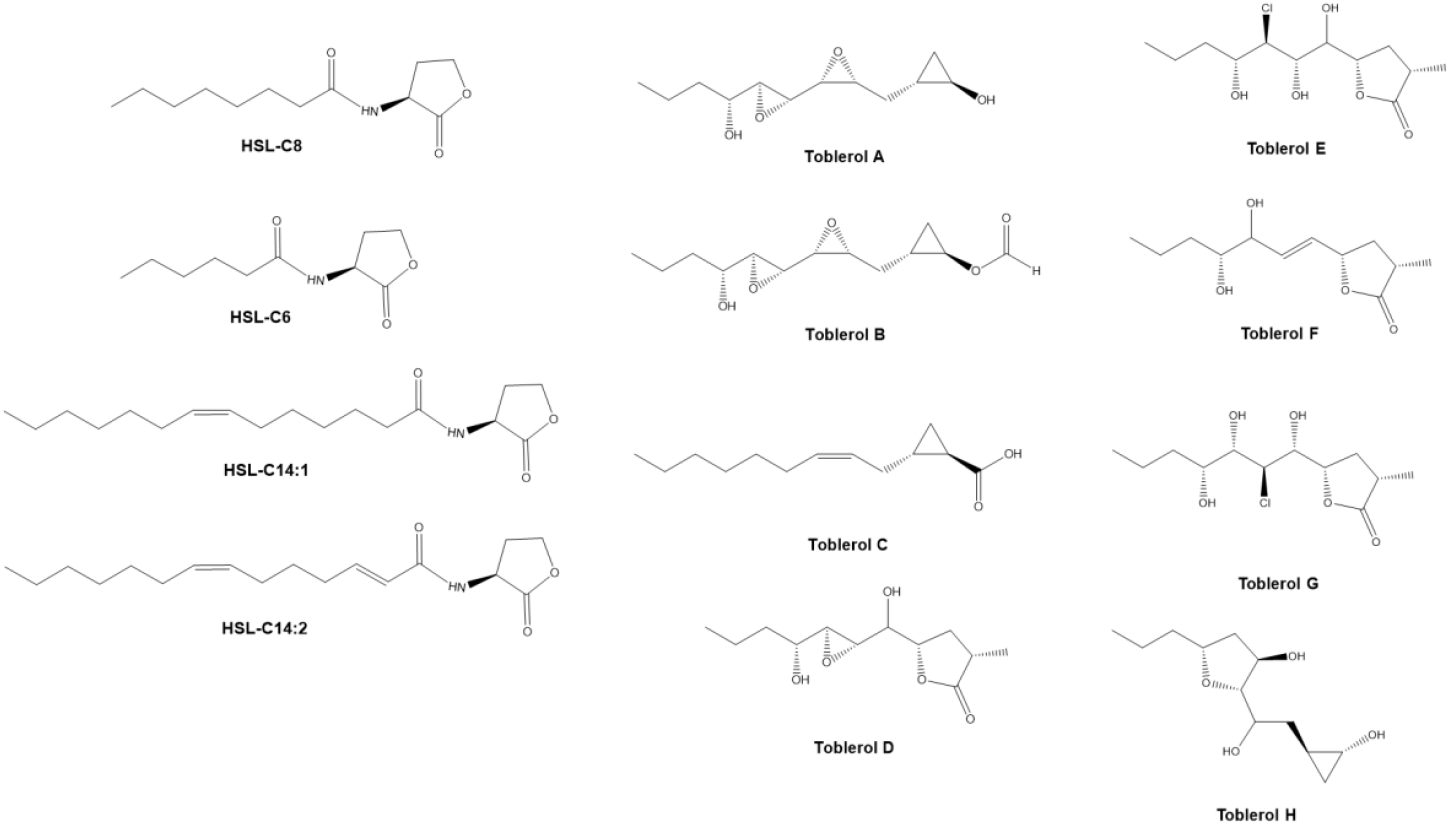
Secondary metabolites isolated from *Methylorubrum extorquens*.

Nowadays, the characterization of known secondary metabolites can be done *in-silico* using genome mining platforms, which look for the association between known biosynthetic gene clusters (BGC) and the consulted genome. A widely used platform is AntiSMASH, which provides the identification, annotation, and analysis of bacterial secondary metabolite biosynthetic gene clusters (Blin et al. 2019; Medema et al. 2011). Parallelly, other techniques to characterize secondary metabolites have been developed, taking advantage of the sensitivity of the current state-of-the-art mass spectrometer instruments. One of these techniques is molecular networking, which associates and clusters molecules based on their MS2 fragmentation pattern (Yang et al. 2013). Molecular networking has facilitated the identification process of known and unknowns molecules (Winnikoff et al. 2014). Recently, molecular networking was also implemented in the characterization of metal-binding molecules (Aron et al. 2022).

The most well-known metal binding molecules are siderophores (Kramer et al. 2020), which are small molecules (up to 2000 Da) that effectively chelate iron(III) by using different chemical functionalities, such as hydroxamates, catecholates, alpha-hydroxycarboxylates, diazeniumdiolate and hybrids of these functionalities (Drechsel et al. 1991; Emery and Neilands 1961; Hardy and Butler 2019; Hermenau et al. 2018; Raymond et al. 2003). The most prolific siderophore producers are bacteria, but siderophores have also been reported in fungi (Ahmed and Holmström 2014).

The discovery of lanthanide dependent microorganisms has presented the question on the potential mechanisms for the acquisition of these metals from the environment (Skovran and Martinez-Gomez 2015). A feasible mechanism that may explain the acquisition of these elements by *M. extorquens* is the production and use of siderophore-like molecules (Daumann 2019; Kramer et al. 2020), which can specifically coordinate, transport and release metals into the cell for their subsequent use as enzymatic cofactors.

Therefore, we herein report the search and characterization of metal chelators from the facultative methylotroph bacterium *Methylorubrum extorquens* AM1. We found the known siderophores N-deoxy-schizokinen A and schizokinen A by using the CAS assay, ^1^H NMR, genome mining, molecular networking and high-resolution mass spectrometry coupled to liquid chromatography. Finally, we measured the capacity of these siderophores to chelate lanthanides by using the arsenazo (III) based assay.

## RESULTS AND DISCUSSION

### Growth and Phylogenetic characterization for *Methylorubrum extorquens* AM1

The strain was cultured in three different liquid culture media, Hypho, MP and Medium 1. Hypho and MP were chosen because they activate methylotrophy in *M. extorquens* AM1, while Medium 1 (M1) is a general culture medium that activates heterotrophic metabolism. All the media produced growth in similar time frames, however the pigmentation produced by Medium 1 was darker than MP and Hypho media (Suppl. Fig. 1). The solid media growth rate showed to be faster in MP and Hypho media than in Medium 1, but this observation can very well be the result of medium adaptation since the strain has been continuously grown in MP and Hypho media.

The phylogenetic characterization of *Methylorubrum extorquens* was performed through the amplification and sequencing of the 16S rRNA gene. We were able to obtain a contig of 1084 nt, which represent around 72% of completeness of the sequence (Suppl. Fig. 2 and 3). Next, we submitted the obtained sequence to NCBI-BLAST to determine the next related type strain. The results showed high percentage of similarity to *M. extorquens* DSM 1337^T^ with a 99.5% of identity. Subsequently, we retrieved 38 sequences of *Methylorubrum* and *Methylobacterium* species from NCBI to determine their phylogenetic placement in consideration with our *in-house* strain, *M. extorquens* AM1 (**Fig 2**). Due to the recent taxonomically emendation of the genus *Methylobacterium* and the transference of eleven of its strains to the new genus *Methylorubrum* (Green and Ardley 2018), we believed that the presentation of the taxonomical characterization is strictly required. The observation of the phylogenetic tree clearly showed that *Methylorubrum* strains generated a different clade than *Methylobacterium* strains, therefore supporting the taxonomical change. In consideration to the working strain, *M. extorquens* AM1, the phylogenetic tree and BLAST analysis provided an identical result, associating *M. extorquens* AM1 with its type strain, *M. extorquens* DSM 1337^T^.

**Figure 2.**
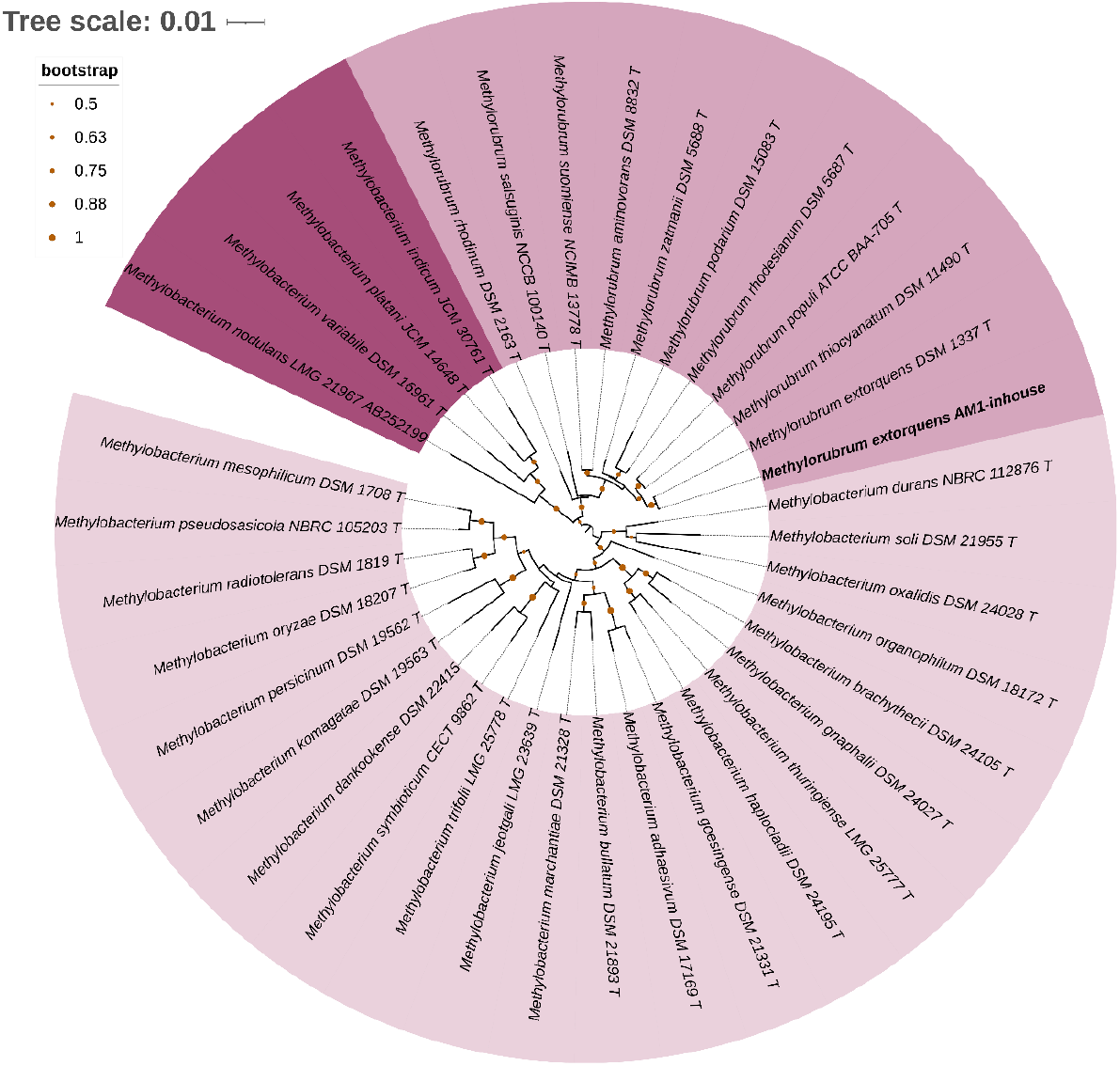
Phylogenetic characterization of *Methylorubrum extorquens* AM1 using a neighbor joining model. Orange dots represent the bootstrap score. The evolutionary distances were computed using the Jukes-Cantor method and are in the units of the number of base substitutions per site (scale). Different colors in the tree are used to characterize the three main clades of the genera *Methylobacterium* and *Methylorubrum*. **T**: type strain. The evolutionary history was inferred using the Neighbor-Joining method. The tree is drawn to scale, with branch lengths in the same units as those of the evolutionary distances used to infer the phylogenetic tree. The evolutionary distances were computed using the Maximum Composite Likelihood method and are in the units of the number of base substitutions per site. This analysis involved 38 nucleotide sequences. All ambiguous positions were removed for each sequence pair (pairwise deletion option). There were a total of 1535 positions in the final dataset. Evolutionary analyses were conducted in MEGA X.

### Genome mining for *Methylorubrum extorquens* and siderophore detection

In general terms, the genetic information of the *Methylorubrum extorquens* AM1 was composed of 5.5 Mb, where the GC content was 69.54%. The coding sequences (CDS) found in the genome represented 5227 genes, of which 73 were pseudo genes.

To investigate the production of secondary metabolites by *Methylorubrum extorquens* AM1, we used the online platform AntiSMASH (Blin et al. 2019). Thus, the concatenated genome and plasmid sequences were submitted and analyzed using relaxed parameters for the search of secondary metabolites. The results of this analysis are shown in **Table 1**.

**Table I.** Found secondary metabolites gene clusters for *Methylorubrum extorquens* AM1. Obtained through AntiSMASH genome mining.

| Location | Secondary metabolites classification | Predicted chemical |
| --- | --- | --- |
| Chromosome | Type I polyketide synthase | NI |
|  | Terpene 1 | Carotenoid |
|  | Siderophore | NI |
|  | Non-ribosomal peptide synthetase 1 | NI |
|  | Terpene 2 | NI |
|  | Homo-serine lactone | Lipopolysaccharide |
|  | Non-ribosomal peptide synthetase 2 | NI |
| Mega-plasmid | Trans-acyl-transferase polyketide synthetase | Toblerols |
|  | Homo-serine lactone | NI |
|  | Bacteriocin | NI |
| Plasmid 1 | Homo-serine lactone | Glycopeptidolipid |
| Plasmid 2 | Not found | - |
| Plasmid 2 | Not found | - |
| *NI: No Information |  |  |

Through the AntiSMASH analysis, eleven different secondary metabolites gene clusters were located and characterized, which were associated to five different classes of natural products enzymes, like polyketides (PKS), non-ribosomal peptides (NRPS), terpene (TP), homo-serine lactones (HSL), ribosomal synthesized and post-translationally modified peptides (RiPPs), which are summarized in **Table I**. The bioinformatic analysis results showed two different gene clusters for PKS production, in which one synthetizes for the recently characterized toblerols, cyclopropanol-containing polyketides (Ueoka et al. 2018b), while the other product is still uncharacterized. AntiSMASH also suggested the production of two NRPS with no published information about their structure and composition. Two different terpenes were also detected, in which one gene cluster showed high association with the biosynthesis of carotenoids. Additionally, AntiSMASH detected in the chromosome and in plasmid 1, the architecture necessary to produce homoserine lactones, which have been already reported in *Methylorubrum extorquens* AM1 (Nieto Penalver et al. 2006). Furthermore, a bacteriocin (RiPPs) with unprecedented architecture was also characterized by the platform. Finally, AntiSMASH found the genetic architecture to produce siderophores (**Figure 3C)**. Interestingly, this same gene cluster was found in multiple *Methylorubrum* and *Methylobacterium* species, as is shown in **Figure 3D**. This strongly suggested that there is a genus-specific production of siderophores in the genera *Methylorubrum* and *Methylobacterium*.

**Figure 3.**
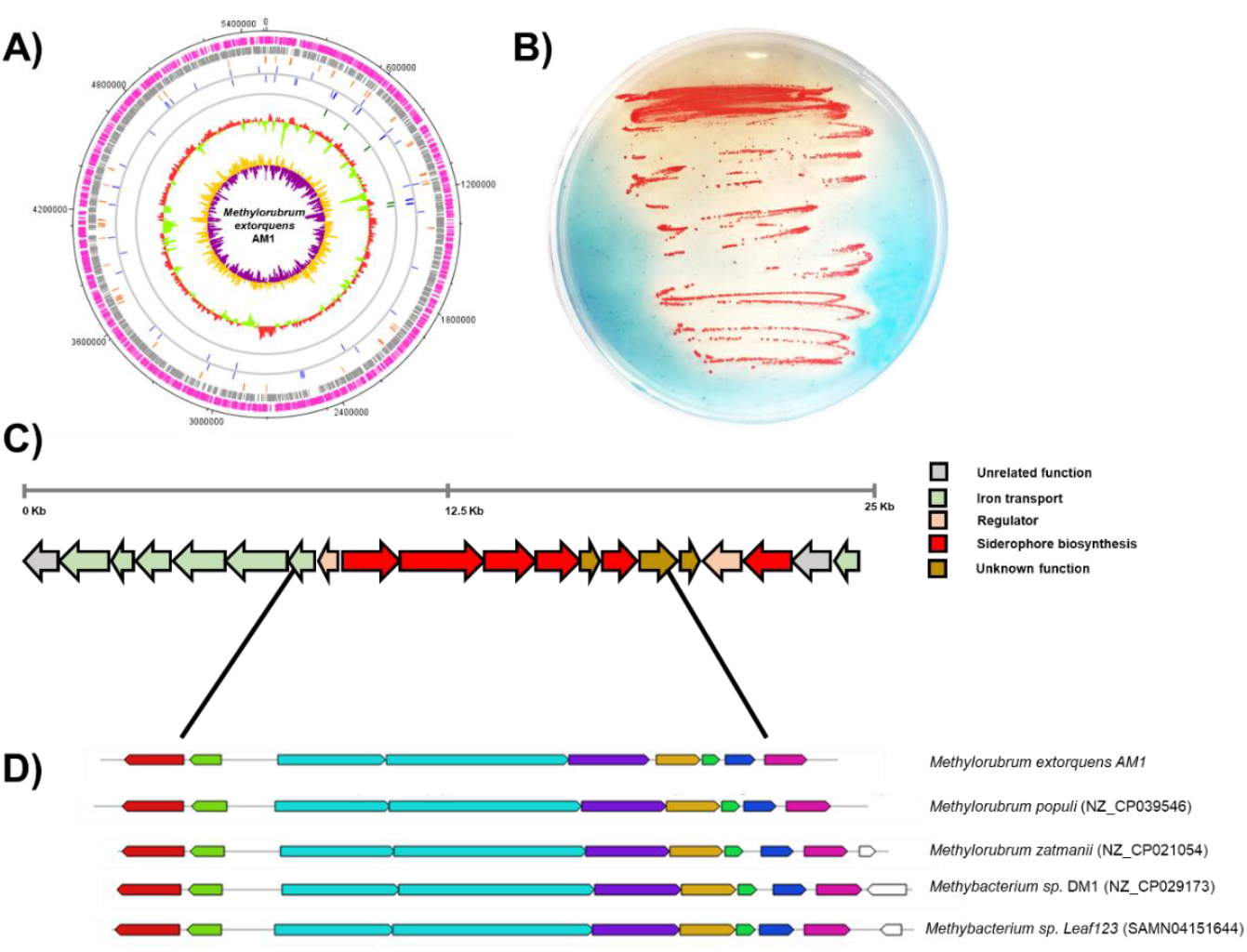
Active siderophore gene cluster contained in the genome *Methylorubrum extorquens* AM1. **3A)** Genome atlas for *Methylorubrum extorquens* AM1. **3B)** CAS-Hypho double layer assay for the detection of siderophores in *Methylorubrum extorquens* AM1. **3C)** Detected *Methylorubrum extorquens* AM1 gene cluster containing iron transport and siderophore production genes. **3D)** Siderophore-core gene cluster found in multiples strains of the *Methylorubrum* and *Methylobacterium* genera. **Genome atlas notation**; Black: DNA base position (bp); Pink: protein-coding regions transcribed in the forward direction; Grey: protein-coding regions transcribed in the reverse direction; Orange: pseudo genes; Blue: tRNA genes; Green: rRNA genes; GC-plot: below average represented by light green, above average represented by red. GC-skew (degree to which the GC content is skewed toward G or skewed toward C): below average represented by purple, above average represented by yellow. The genome plot was generated using DNAPlotter version 1.4 from Artemis (Sanger Institute).

Since our interest was the search of chemicals that can effectively coordinate with metals, we subsequently analyzed the core of the putative siderophore biosynthetic gene cluster (BGC) to discern which was the most similar BGC and its product. This latter evaluation indicated that the genetic information in *Methylorubrum extorquens* AM1 had no high homology to other known gene clusters. The highest score that was obtained was in the range of 30-39% similarity and it pointed to petrobactin, a bis-catecholate-α-hydroxy acid siderophore. However, this score is weak, thus this latter siderophore couldn’t be considered as a potential candidate for *M. extorquens* AM1.

To determine if the siderophore gene cluster was active in the strain, we performed the a modified version of the CAS assay (Louden et al. 2011) using a double layered method, which allowed us to bypass the potential toxicity the CAS reagents could produce on *Methylorubrum extorquens* AM1. The results of this qualitative analysis are shown in **Fig 3B**, where it can be observed that *M. extorquens* AM1 indeed was able to produce siderophores, which captured the iron from the CAS reagent and produced the coloration shift (from blue to light brown), characteristic for siderophore production. As the genome mining analysis was not able to reveal the identity of the siderophore being produced, we undertook the chemical analysis of *Methylorubrum extorquens* AM1.

### Growth scale up, chemical extraction and profiling

To determine the identity of the siderophores being produced by *Methylorubrum extorquens* AM1, we undertook growth experiments using three different cultures media. Two of these media (Hypho and MP) activated methylotroph metabolism, meanwhile Medium 1 activated heterotrophic metabolism. After 13 days of incubation, we harvested the culture and proceeded with the chemical extraction. The extraction was identical for each media, and used amberlite XAD-16. After the secondary metabolites extraction, we obtained the extraction yields, which showed that Medium 1 produced 25 mg/L of crude extract. In contrast, medium Hypho only produced 13.5 mg/L, which was very similar to the yield produced by medium MP that reached 15 mg/L. Subsequently, we proceeded with the analysis of the samples through LCMS to obtain the chemical profiles of the samples. From this last analysis, we observed that the chemical profiles at 210 nm for medium Hypho and MP were almost identical (Suppl. Fig. 4), which is reasonable since both media triggered methylotrophic metabolism. On the other hand, Medium 1 produced a UV 210 nm profile that differed from Hypho and MP media, being less complex at first sight. This latter observation resulted to be interesting, since it is common knowledge that heterotrophic metabolism can use a larger variety of nutrients, which in turn, should activate a larger number of metabolic pathways and their products. As we observed that Hypho and MP medium produced quite similar results, we continued with only one methylotrophic medium, in this case, Hypho medium because some metabolites had higher peak intensities in comparison with MP medium (Suppl. Fig. 4).

To gain further insights from the samples, we characterized the ^1^H NMR profile of the crude extracts for the Hypho and M1 media (Suppl. Fig. 5). From the ^1^H NMR spectra, it can be inferred that the amberlite extraction resulted in a complex mix of metabolites, which differ in their chemical group functionalities (chemical shift function). This last statement confirmed that the extraction strategy was working properly regarding the chemical diversity. Interestingly, exchangeable protons were observed in the high frequency spectrum zone, which may be associated to the tautomerization processes of carboxylic acids and neighboring ketone-hydroxyl proton exchange. Additionally, it was possible to observe a few singlet-broad peaks that can be associated with nitrogen-related functionalities. Moreover, several signals for olefinic, aromatic and alkyl groups were observed in both samples, as well as the characteristic signals for cyclopropane (<1 ppm), which belong to toblerol derivatives (Ueoka et al. 2018a). Finally, when both samples were compared side by side, it was clear that the crude extract of the methylotroph medium, Hypho, had a larger chemical diversity than M1 (Suppl. Fig. 5).

Subsequently, we undertook the mass spectrometry analysis of the samples. As we recognized the complexity of the crude extracts from the HPLC and NMR analysis, we had to select a feasible method to analyze large datasets. Hence, we selected the Global Natural Products Social Molecular Networking, GNPS (Yang et al. 2013), which allowed the visualization of the entire mass spec dataset for its analysis.

### Molecular networking and siderophore identification

The crude extracts produced by the growth of *Methylorubrum extorquens* AM1 in two culture media (Hypho and Medium 1) were analyzed through high pressure liquid chromatography coupled to a qTOF mass spectrometer. Next, from the acquired data we built the molecular networking of both samples in parallel by using the MS2 mass spectrometry data. We selected this method because it allows the analysis of the large dataset in a single view (Suppl. Fig. 6). The analysis of the molecular networks showed a quite complex structure, where both media produced similar results. Through the molecular networking visualization and the complementation with the LCMS data, we could determine the production of several known *Methylorubrum extorquens* AM1 secondary metabolites. For instance, we observed the production of the toblerols series in both culture media. A similar situation was found regarding the known homoserine lactones produced by *M. extorquens* AM1. Interestingly, the strain in both culture media was able to produce > 50 chemical clusters which can be translated to a large chemical diversity involving secondary metabolites. Usually, Gram-negative bacteria are thought to be low producers of secondary metabolites in comparison with Gram-positive strains, ex. *Streptomyces* strains. In our particular case, we compared the molecular networking produced by *Streptomyces cacaoi* (Liu et al. 2020) against *M. extorquens* AM1 and we observed that both networks shared a high number of chemical clusters, providing evidence that Gram-negative bacteria can also be a prolific source of secondary metabolites with potential for biotechnological application.

In our search for a metal chelator from *M. extorquens* AM1, we paid attention to a small cluster that was present in the molecular networking (**Fig 4A**), where we found a m/z 387.18 and m/z 404.20. The former m/z corresponded to the [M+H]^+^ ion and the latter [M+H_2_O]^+^ ion, which helped to identify the exact mass of the target. Then, with the exact mass, we calculated the following molecular formula, C_16_H_26_ N_4_O_7_, which was in agreement with the molecular formula of N-deoxyschizokinen A. N-deoxyschizokinen A is a citrate-hydroxamate molecule which derives from the schizokinen series siderophores (Chuljerm et al. 2020; Mullis et al. 1971). To confirm that our finding was correct, we analyzed the LCMS data manually, determining that indeed N-deoxyschizokinen A was present in the sample (**Fig. 4-B2**).

**Figure 4.**
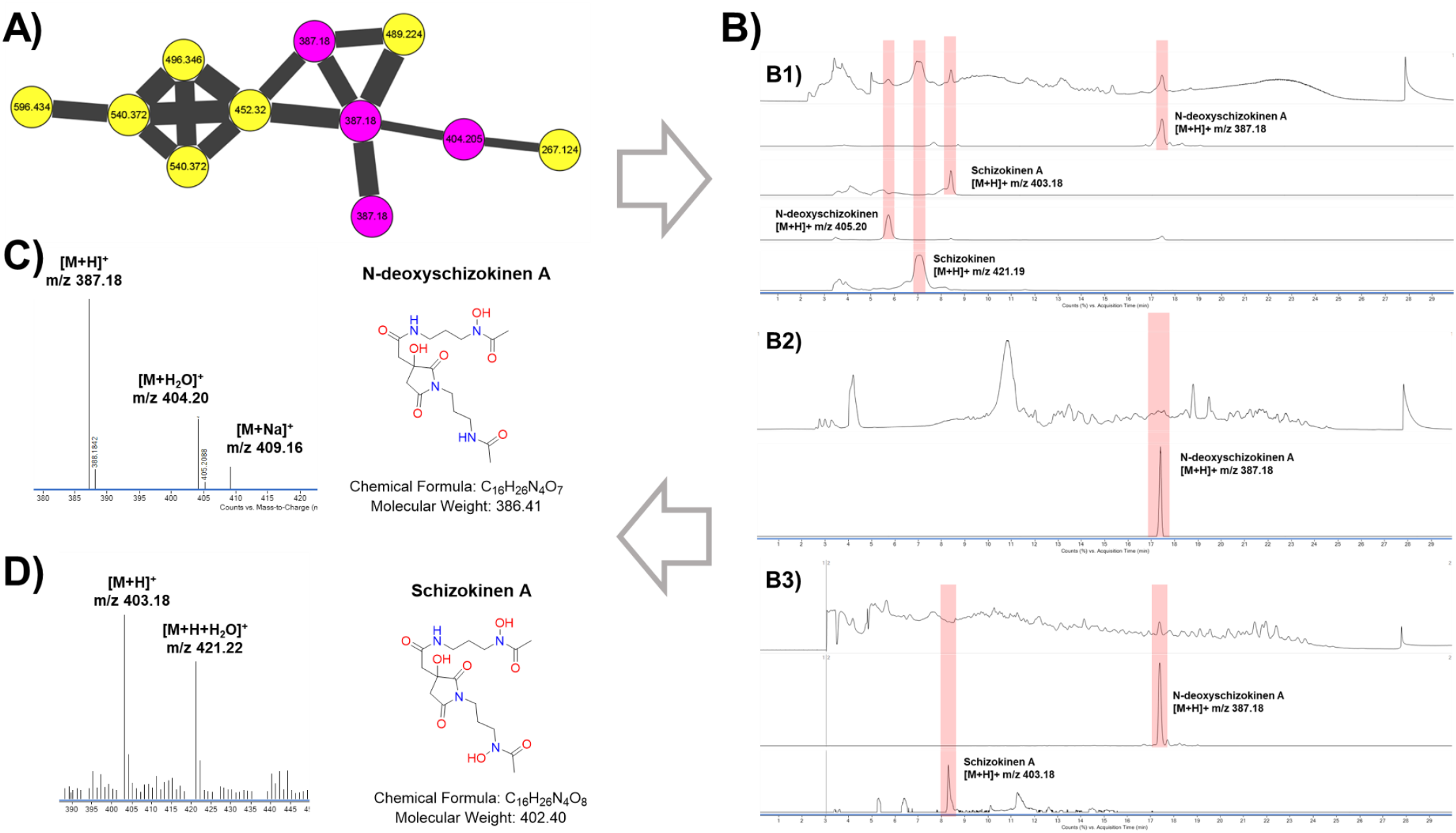
Identification of the siderophores N-deoxyschizokinen A and schizokinen A from *Methylorubrum extorquens* AM1. **4A)** Molecular networking cluster showing the [M+H]^+^ and [M+H_2_O]^+^ ions for N-deoxyschizokinen A. **4B-1)** Top chromatogram: TIC of the schizokinen series standard, lower chromatograms: EIC for each schizokinen derivative. **4B-2)** Top chromatogram: TIC for the amberlite XAD-16 extraction, bottom chromatogram: EIC for N-deoxyschizokinen A. **4B-3)** op chromatogram: TIC for the Phenol-Chloroform extraction, middle chromatogram: EIC for N-deoxyschizokinen A, bottom chromatogram: EIC for schizokinen A. **C)** Mass spectrum and chemical structure for N-deoxyschizokinen A. **4D)** Mass spectrum and chemical structure for schizokinen A.

Finding a single derivative of the schizokinen series was intriguing because they have been reported in tandem (Hu and Boyer 1995). Therefore, we assumed that our resin base amberlite XAD-16 extraction was not effective to obtain the complete chemical series. Thus, to determine if the production of other schizokinen series molecules was possible, we undertook a second chemical extraction, this time using an old fashion siderophore extraction method that involved a phenol-chloroform mix. This same methodology was used for the first schizokinen publication (Mullis et al. 1971). The results for this extraction are presented in **Fig 4-B3**, where it can be observed that a second schizokinen derivative could now be detected. Despite our chemical efforts, schizokinen and N-deoxyschizokinen were not detected in *M. extorquens* AM1 cultures. It has been reported in different studies (Hu and Boyer 1995; Mullis et al. 1971) that schizokinen and N-deoxyschizokinen undergo chemical decomposition by heat or acidic conditions to produce the cyclic imide derivatives, schizokinen A and N-deoxy-schizokinen A. Therefore, it is reasonable to think that the whole schizokinen series (**Fig. 5**) was being produced by *M. extorquens* AM1, but under the growing or chemical extraction conditions both molecules, schizokinen and N-deoxy-schizokinen, were totally transformed to their cyclic imide derivatives, schizokinen A and N-deoxy-schizokinen A (**Fig 4C-D**).

**Figure 5.**
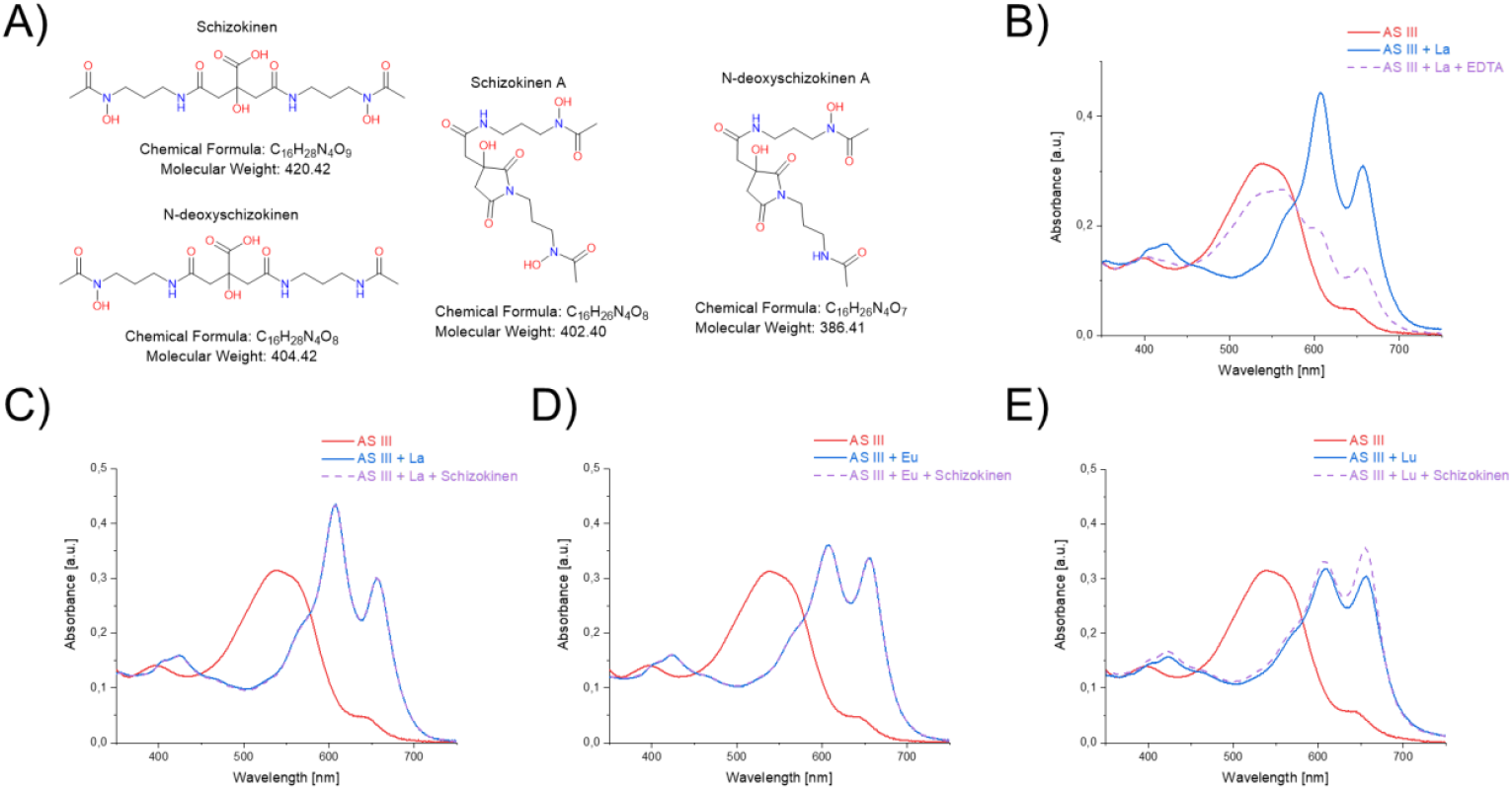
Arsenazo (III) assay to determine lanthanide chelation process of the schizokinen series siderophores. **5A)** Chemical structures for all the members of the schizokinen series. **5B)** Spectral results for the arsenazo-lanthanum complex and EDTA, which was the positive control. **5C)** Spectral results for the arsenazo-Lanthanum complex and the schizokinen siderophores. **5D)** Spectral results for the arsenazo-Europium complex and the schizokinen siderophores. **5E)** Spectral results for the arsenazo-Lutetium complex and the schizokinen siderophores.

Next, to add one more layer of certainty to our findings, we obtained a schizokinen series standard, which contained schizokinen, schizokinen A, N-deoxyschizokinen and N-deoxyschizokinen A (**Fig 5A**) and we analyzed it through LCMS to compare our results in consideration to retention time and molecular weight. The results for the schizokinen standard results are presented in **Fig. 4-B1**. By comparing **Fig. 4B-1, Fig. 4B-2 and Fig. 4B-3**, it is evident that the two molecules detected by our experiments, schizokinen A and N-deoxyschizokinen, indeed showed identical chemical characteristic as the schizokinen series standard. Finally, the mass spectra and the chemical structures of the detected schizokinen derivatives are shown in **Fig. 4-C and Fig. 4-D**.

A recently published work by our collaborators in the University of California at Berkeley (Zytnick et al. 2022), reported that *M. extorquens* AM1 highly upregulated the siderophore gene cluster when grown using a methylotroph culture medium (Hypho) supplemented with Nd_2_O_3_, thus showing a direct relationship between siderophore production, methylotrophy and rare-earth metals. To determine if this finding was also true for our study, we compared the relative concentration of N-deoxyschizokinen A in a methylotrophic and heterotrophic culture media both supplemented with the rare-earth metal, Lanthanum. Both growing conditions, Hypho and Medium 1, were incubated with identical conditions to determine which was able to produce the largest quantity of N-deoxy-schizokinen A. As it can be seen in Suppl. Fig. 7, Hypho medium is considerably a better condition to produce the schizokinen derivative, because its biosynthesis is increased by at least nine times. This last finding also confirmed a relationship between methylotrophy and metal chelation.

### Lanthanide chelation activity

The found relationship between methylotrophy, rare-earth metals and siderophores opened the possibility that the found molecules could chelate lanthanides. To date, schizokinen derivatives are known metals chelators, which have been shown to chelate; aluminum, copper, and iron (Arceneaux et al. 1984; Hu and Boyer 1996; Mullis et al. 1971). Therefore, it would be reasonable to think that schizokinen derivatives have certain plasticity to chelate different metals. Following this premise, we undertook the analysis and determination of the lanthanide-chelation activity of this siderophore family (**Fig. 5-A**). Thus, we took advantage of the facile arsenazo III-Based Assay (Hogendoorn et al. 2018) and used EDTA as positive control (**Fig. 5-B**). The facile arsenazo (III) based assay relies in the absorbance shift produced by the subtraction of the lanthanide element from the metal-arsenazo (III) complex, which in turn produces a coloration change that can be determined by a spectrophotometer. Subsequently, we proceeded with the measurements using three different lanthanides, Lanthanum (early), Europium (mid-point) and Lutetium (heavy). The results are presented in **Fig. 5C-D**. As can be observed in the charts, the schizokinen siderophore series did not show any chelation activity on the tested lanthanides even when the whole series was used at the same time. The result nonetheless was not surprising, because the schizokinen series siderophores (individually or grouped) doesn’t satisfy the coordination number required for rare-earth metal chelation, where the minimal required coordination number for lanthanides is larger than 6 (Cotton 2005; Lumpe et al. 2020). Therefore, this finding suggested that the schizokinen series siderophores were not involved in lanthanides chelation, transport, and delivery.

Recently, a *Streptomyces* derived polyether ionophore, monensin, showed the ability to complex lanthanum (La^3+^) and neodymium (Nd^+3^) after the deprotonation of the carboxylic acid moiety (Pantcheva et al. 2019). It would be interesting to see if other polyether molecules as maitotoxin, gymnocin A, brevetoxin B that are produced by marine microorganisms (Nicolaou et al. 2008) can have the capacity to coordinate lanthanides as monensin does. The experimentation with aza-crown ether complexes has also shown to coordinate lanthanides (Itoh et al. 2000). In the case of a generalized lanthanide chelation activity of these polyethers, we would be in the presence of the first case of a nature-made small molecule capable of chelating and transporting lanthanides, which can open the doors to new mining technologies for rare-earth metals.

## CONCLUSION

Through the bioinformatic and microbiological analysis of the Gram-negative and lanthanide user bacterium *Methylorubrum extorquens* AM1, we determined the presence of a biosynthetic gene cluster for the production of siderophores. The direct observation of *M. extorquens* AM1 cultures on the CAS assay indicated the functionality and production of siderophores by the gene cluster. Subsequently, by different culture conditions and chemical extractions of *M. extorquens* AM1, we determined the production of two known siderophores, schizokinen A and N-deoxyschizokinen A. Interestingly, the other two known siderophores of the series, schizokinen and N-deoxyschizokinen, could not be detected in our cultures. Very likely, our means of extraction produced the total transformation from the open to the cyclic form of the siderophores. After the characterization of the molecules, we proceeded to determine the lanthanide chelation activity by the facile arsenazo (III) assay as *M. extorquens* AM1 is a known rare-earth metals user. Our studies showed that the schizokinen siderophores do not have any chelation activity to Lanthanum, Europium and Lutetium, which rules out their involvement in the lanthanide acquisition process by *M. extorquens* AM1. Finally, further research needs to be performed to determine if *M. extorquens* AM1 uses a siderophore-like molecule for the acquisition of rare-earth metals from the environment.

## METHODOLOGY

### Cultivation

Three different media were used for the cultivation of *Methylorubrum extorquens* AM1, MP, Hypho and Medium 1 (M1). **MP medium** (for 1L: sodium citrate (Na_3_C_6_H_5_O_7_*2H_2_O) 0.0134 g, PIPES (C_8_H_8_N_2_O_6_S_2_) 9.07 g, K_2_HPO_4_*3H_2_O 0.33 g, NaH_2_PO_4_*H_2_O 0.259 g, MgCl_2_*6H_2_O 0.1 g, (NH_4_)_2_SO_4_ 1.057 g, CaCl_2_*2H_2_O 2.94 mg. The trace elements composition was: ZnSO_4_*7H_2_O 0.345 mg, MnCl_2_*4H_2_O 0.19 mg, FeSo_4_*7H_2_O 5 mg, (NH_4_)_6_Mo_7_O_24_*4H_2_O 2.4 mg,CuSO_4_*5H_2_O 0.25 mg, CoCl_2_*6H_2_O 0.475 mg, Na_2_WO_4_*2H_2_O 0.1 mg. The pH of the medium was adjusted to 7.3), **Hypho medium** (for 1L: K_2_HPO_4_*3H_2_O 3.309 g, NaH_2_PO_4_ 2.255 g, MgSO_4_*7H_2_O 0.1972, (NH_4_)_2_SO_4_ 0.5 g. All dissolved in 1L of distillated water. The trace elements composition was based in a modified version of Vishniac solution (Vishniac and Santer 1957). 3 mL/L were added of solution A (FeSO_4_*7 H_2_O 0.1 g, EDTA 0.7849 g, dissolved in 300 mL of distillated water with the addition of concentrated NaOH). 2.5 mL/L were added of solution B (CaCl_2_*2H_2_O 0.14 g, MnCl_2_*4H_2_O 0.1 g, (NH_4_)_6_Mo_7_O_24_*4H_2_O 0.02 g, CuSO_4_ 0.03 g, CoCl_2_*6H_2_O 0.032 g, ZnSO_4_*7H_2_O 0.44 g. Dissolved in 250 mL of distillated water). The pH of the medium was adjusted to 7.1) and **Medium 1** (for 1L: peptone 5 g, meat extract 3 g, pH adjusted to 7.0). All the culture media were supplemented with 5.5 mL of MeOH and a final concentration of 2 µM of LaCl_3_*7H_2_O. The culture media was autoclaved at 121°C during 30 min. For solid culture media, we added Agar 15 g/L. The incubation time for the liquid cultures was 13 days. The culture conditions for the liquid cultivation were as following; 29°C, 250 RPM, daylight cycle, Ultra-Yield® Flasks (Thomson, San Diego, California, USA). The inoculation process was made by the addition of 1 cm^2^ of a solid *M. extorquens* AM1 culture (7 days old), which was inoculated into a 250 mL flask of the selected medium to produce the seed culture (5-8 days). Subsequently, 25 mL of the seed culture were inoculated into the 2.5 L growth flask. After 13 days, the culture was harvested and extracted chemically. The conservation of *Methylorubrum extorquens* AM1 was performed by selecting an axenic culture, from which, a single pure colony was taken and transferred to the cryopreservation system of MAST CRYOBANK® (Mast Group, Bootle, UK.) following the manufacturer instructions. The conserved cells were stored at -20°C.

### Bioinformatics

The genome of *Methylorubrum extorquens* AM1 is available publicly (NCBI access # ASM2268v1). The files contained the genome and plasmid sequences of *Methylorubrum extorquens* AM1 in FASTA format. To visualize the genome and plasmid sequences, a genome atlas was constructed using DNAPlotter version 1.4 from Artemis (Carver et al. 2011). This latter software was also used for genome screening. The genome mining of *Methylorubrum extorquens* AM1 was developed with AntiSMASH, which allows the rapid genome-wide identification, annotation and analysis of secondary metabolites biosynthesis gene clusters in bacterial genomes (Blin et al. 2019). Protein homology of selected regions of the genome of *Methylorubrum extorquens* AM1 were evaluated through the NCBI tools-BLAST (Altschul et al. 1990).

### Molecular Characterization

The molecular characterization of *Methylorubrum extorquens* AM1 was prepared by taking a single colony from an axenic culture and the subsequent DNA extraction using DNeasy Qiagen™ kit (Qiagen GmbH, Hilden, Germany). For the amplification of the 16S rRNA gene sequence, we used PCR and the following primers: 27F, 1492R, 534R and 342F, 10 µmol/µL (Lane 1991). The reagents for PCR were obtained from GE Healthcare illustra™ PuReTaq Ready-To-Go™ PCR Beads containing DNA polymerase, MgCl_2_, and dNTPs (GE Healthcare, Glattbrugg, Switzerland). Each reaction tube contained 1 µL of primer, 19 µL of DNA free water, 5 µL of the DNA solution. The sequencing process was run by the company Eurofins (Germany) through its Sanger sequencing GATC. The 16S rRNA gene sequences were manually curated and assembled using Chromas pro software (version 1.7.6) and saved in FASTA format. Sequences were first identified with the Ribosomal database project (rdp.cme.msu.edu) (Cole et al. 2013). The alignment of the sequences was performed by using BLAST (Altschul et al. 1990). The parameter used with BLAST was type strain.

### Phylogenetic Analysis

For the phylogenetic tree constructions, the closest reference strains were downloaded from the NCBI platform. The placement of the reference strains was compared with those of the ARB project (Ludwig et al. 2004; Westram et al. 2011). Next, the sequences were aligned using SILVA web tool SINA (Quast et al. 2013). The alignment was saved in FASTA format. MEGA X software (version 10.2.4) was used to delete gaps and to calculate phylogenetic models (Tamura et al. 2011). Based on the results of the MEGA X model evaluation, we accomplished bootstrapped phylogenetic trees using the maximum likelihood and neighbor joining models. The final construction of the phylogenetic trees was made through the use of the iTOL web platform (Letunic and Bork 2021).

### Chemical extraction of *Methylorubrum extorquens* AM1

The chemical extraction was performed after the cultivation phase is finished. The cultures were centrifuged at 8000 RPM, 20°C, during 15 min. Then, the supernatant was separated from the cells. Next, 20 g/L Amberlite XAD-16 (Alfa Aesar, Kandel, Germany) were added to 1L of supernatant and incubated for 1 h. Subsequently, Amberlite XAD-16 was recovered from the supernatant, washed with 2 L of distillated water to be finally eluted with acetone (HPLC grade, Sigma-Aldrich, St. Louis, Mo USA). This last elution is brought to dryness using a rotational evaporator under reduced pressure. Then, the dried acetone residue was resuspended in distillated water and placed in a separation funnel, where it was extracted 3 × 300 mL ethyl acetate (HPLC grade, Sigma-Aldrich, St. Louis, Mo USA). The organic phase was recovered, concentrated until dryness, and stored in a glass vial at -20°C for its subsequent use. The second extraction methodology was based on the phenol-chloroform (1:1) extraction for siderophores and we proceeded as reported in Mullis et at. (Mullis et al. 1971).

### ^1^H Nuclear magnetic resonance analysis

^1^H NMR profiles of *Methylorubrum extorquens* AM1 both growing conditions, M1 and Hypho media were recorded by dissolving the crude extracts in CDCl_3_ (Eurisotop, Saint-Aubin, France) supplemented with CD_3_OD (Deutero GmbH, Kastellaun, Germany) drops. Then, the samples were transferred to NMR tubes (regular and Shigemi tubes) and acquired in a 600 MHz Bruker instrument. TMS (Eurisotop, Saint-Aubin, France) was added as internal standard. The NMR data was processed by Mnova software, as well as with Bruker TopSpin.

### Detection, HRLCMS-qTOF analysis and mass networking

The separation and determination of the molecular weight and exact mass of the molecules was performed in a 1260 Agilent Infinity II HPLC instrument coupled with Agilent 6530 Q-TOF LC-MS mass spectrometer. The detection of the molecules was performed by three means. Using the diode array detector (DAD) at 210 nm of an Agilent 1260 Infinity II HPLC system (Agilent Technologies Inc., Santa Clara, California, USA), total ion current (TIC) chromatogram and the extracted-ion chromatogram (EIC) by using an Agilent 6530 Q-TOF LC/MS mass spectrometer (Agilent Technologies Inc., Santa Clara, California, USA). For the analysis of schizokinen derivatives in the crude extracts and the original standard, we used a 30-minute gradient of water and acetonitrile supplemented with 0.1% LCMS grade formic acid (Fisher chemical, Geel, Belgium), and employing an Agilent 1260 Infinity II HPLC system (Agilent Technologies Inc., Santa Clara, California, USA). The chromatographic column used was NUCLEODUR C18 Isis, 5 µm, 250 × 4.6 mm from Macherey-Nagel (Dueren, Germany). The gradient used was as follows: 0 min 90% A 10% B, 2 min 90% A 10% B, 20 min 0% A 100% B, 23 min 0% A 100% B, 28 min 90% A 10% B, 30 min 90% A 10% B, where A is LCMS grade water (Merck, Darmstadt, Germany) and B: LCMS grade acetonitrile (VWR, Fontenay-sous-Bois, France) both supplemented with 0.1% LCMS grade formic acid (Fisher chemical, Geel, Belgium). The flow rate used was 0.7 mL/min. The instrument was controlled through MassHunter (Agilent Technologies Inc., Santa Clara, California, USA). The mass spectrometer was equipped with a Jet Stream Technology Ion Source (Agilent Technologies Inc., Santa Clara, California, USA). The parameters applied for the determination of the exact mass were as following: positive mode, gas temperature: 350°C, drying gas flow: 10 L/min, nebulizer pressure: 45 psi, sheath gas temperature: 400°C, sheath gas flow: 12L/min, VCap: 3500 V, nozzle voltage: 0 V, fragmentor: 100 V, skimmer: 65 V, oct RD Vpp: 750 V. The mass range used for the experiments was 50-1000 Da. MassHunter software was used to process the data. The injection volume for each sample was 15 µL, the flow rate was 0.7 mL/min. The crude extract samples were injected in concentration equal to 0.5 mg/L. To identify the molecules from the schizokinen standard or *Methylorubrum extorquens* AM1 crude extracts, we used the extracted ion chromatogram (EIC) in conjunction with the schizokinen derivatives exact masses. N-deoxyschizokinen A [M+H]^+^: 387.18, schizokinen A [M+H]^+^:403.18, N-deoxyschizokinen [M+H]^+^: 405.19 and schizokinen [M+H]^+^:421.19. The samples were filtered through PTFE 0.22 µm filter (Fisher scientific, China) prior to their injection. After the injection, the samples were stored at -20°C

For the mass networking data acquisition, we used the exact methodology and instrument that in the MS1 analysis, but this time acquiring MSMS (MS2) data instead of MS1. The parameters used for the mass-tandem fragmentation experiments were as following: positive mode, gas temperature: 350°C, drying gas flow: 10 L/min, nebulizer pressure: 45 psi, sheath gas temperature: 400°C, sheath gas flow: 12L/min, VCap: 3500 V, nozzle voltage: 0 V, fragmentor: 100 V, skimmer: 65 V, oct RD Vpp: 750 V. The mass range used for the experiments was 50-2000 Da. The fragmentation energy was done through a ramped collision energy function, where the slope was 6 and the offset 4. For the construction of the molecular networking, we followed the procedure indicated by Yang et al. (Yang et al. 2013). Next, the data was analyzed and curated for accuracy. Subsequently, the data was submitted to the GNPS platform, and the network constructed. The visualization of the network was done using Cytoscape (Version: 3.8.2).

### Schizokinen siderophore standard

Schizokinen, deoxyschizokinen, schizokinen A and deoxyschizokinen A were obtained from Biophore Research Products (Tübingen University, Tübingen, Germany) as a single mix of schizokinen derivatives. Schizokinen and its derivatives were dissolved in a mixture of methanol and water (1:1). The injection concentration and volume for the standard solution was 0.1 mg/mL and 50 µL. The samples were filtered through PTFE 0.22 µm filter (Fisher scientific, China) prior to their injection. After the injection, the samples were stored at -20°C.

### CAS and Arsenazo(III) assays

We performed the CAS assay (Schwyn and Neilands 1987) by modifying the procedure described by Louden et al. (Louden et al. 2011). Therefore, we made a double layer CAS assay, where the CAS solution was placed on the bottom of a 15 cm Petri dish and on top of it, we poured solid Hypho culture medium (see Cultivation section). Both solutions, CAS and Hypho, were autoclaved at 121°C before being poured in the Petri dishes. Once the plates were cooled down, we stored them in darkness until their subsequent use. The inoculation of the plates was performed by using a metallic loop and *Methylorubrum extorquens* AM1 axenic cultures. The plates were incubated at room temperature for 30 days in darkness.

The Arsenazo (III) assay was performed based on the protocol described by Hogendoorn et al (Hogendoorn et al. 2018). 1 mM AS III stock solution was prepared in MilliQ water in an amber vessel, and then diluted to a final concentration of 200 µM. Similarly, 200 µM stocks of LaCl_3_*7H_2_O, EuCl_3_*6H_2_O and LuCl_3_*6H_2_O stocks were prepared in MilliQ water. A single 200 µM solution of all the schizokinen siderophores was also prepared in MilliQ water. A two-component buffer containing 5 mM MOPSO and 5 mM CHES, pH=6, was used for the measurements. The measurements were performed stepwise, where step one was to collect only the spectrum of a 10 µM AS(III) in the buffer solution. Then, an equimolar amount of the lanthanide was added, generating another read of the spectrum profile. Next, 10 µM of the mix of schizokinen siderophores was added, and the spectra were collected again.

The positive chelation control was a solution of EDTA. The 200 µM EDTA solution was added to an AS(III)-lanthanides complex. Subsequently, the spectra were recorded. All the measurements were performed in a Cary 60 spectrophotometer (Agilent Technologies Inc., Santa Clara, California, USA) using single-use cuvettes with a pathlength of 10 mm. A baseline correction was performed prior to each measurement, and the spectra were corrected to dilution.

## ACKNOWLEGMENTS

IS is thankful for the support provided by the Chemistry faculty of LMU and to the dean, Prof. Dr. Philip Tinnefeld. IS thanks Helena Singer for her support with the Arsenazo experiments.

## SUPPORTING INFORMATION AND DATA AVAILABILITY

Supporting information and complementary data is available in the supplementary information file.

